## Supplementary Figures 1-6 for "Circular RNAs in the human brain are tailored to neuron identity and neuropsychiatric disease"

### Supplementary Fig. 1

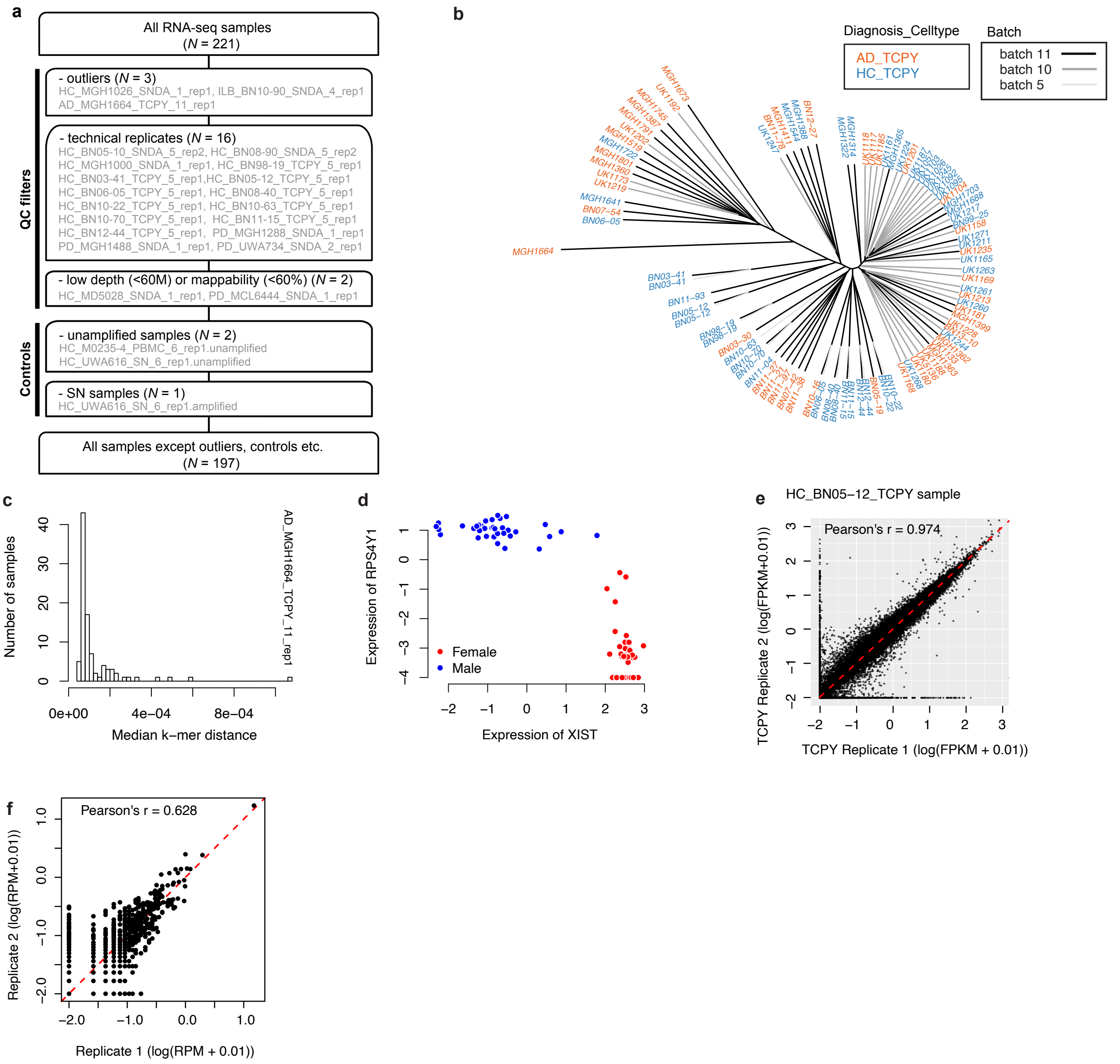


**Supplementary Figure 1. RNA-seq sample filters and quality control, including outlier detection, sex concordance, and assay performance measures**. **a**, Schematic of RNA-seq sample filtering. IDs of excluded samples are listed under each filtering step in gray text. Two QC tests were performed to identify outlier samples based on systematic abnormalities in overall expression (**b**,**c**). Moreover, we tested for sex concordance to identify potential sample mix-ups (**d**). **b**, Dendrogram visualizing pairwise Spearman correlations between gene expression levels of temporal cortex neuronal samples that are newly added after our previous study (Dong et al. Nature Neuroscience, 2018). **c**, Histogram of median pairwise k-mer distances for each of the 221 samples with all other samples. **d**, Concordance between clinical sex and sex-specific gene expression in neuronal and non-neuronal samples: normalized expression levels of the female-specific *XIST* transcript (x axis) and normalized expression levels of the Y-chromosome specific *RPS4Y1* transcript (y axis) are shown. **e**, Scatterplot of two technical replicates based on lcRNAseq. N = 57,814; all annotated genes in GENCODE v19. **f**, Scatterplot of circRNA expression from two technical replicates (panel e) based on lcRNAseq.

### Supplementary Fig. 2

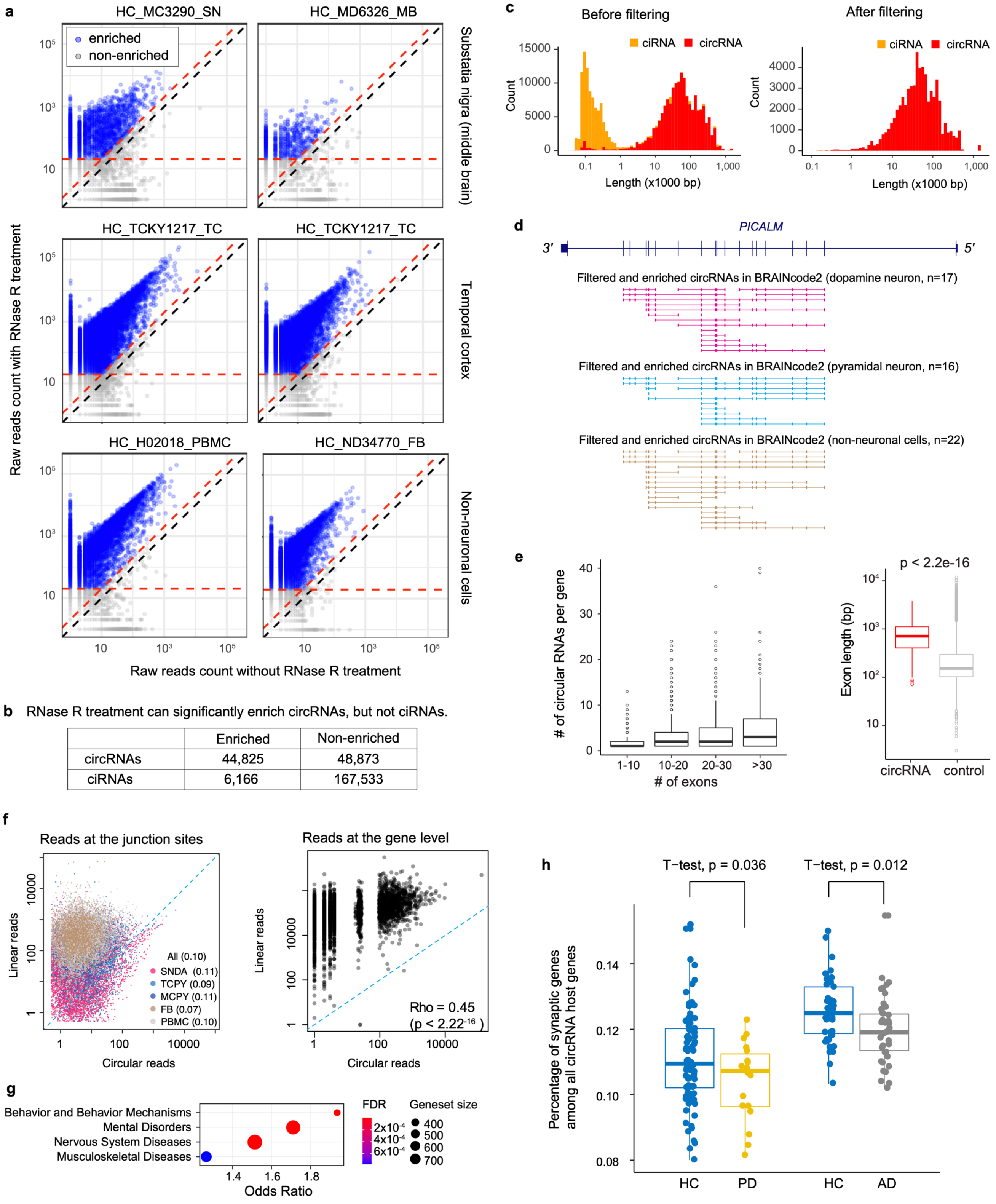


| | Enriched | Non-enriched |
|---|---|---|
| circRNAs | 44,825 | 48,873 |
| ciRNAs | 6,166 | 167,533 |

**Supplementary Figure 2. Circular RNA charateristics**. **a**, Scatterplot of circRNA reads count between paired RNase R-treated vs. mock-treated RNA-seq samples. Blue dots are those enriched in RNase-R treatment based on (1) ratio between RNase treated vs. mock reads is greater than or equal to 2, and (2) at least 20 reads in the RNase treated RNA-seq. **b**, Number of circRNAs (exon-derived) and ciRNAs (intron-derived) enriched by RNase R treatment. It showed that RNase R treatment can significantly enrich exon-derived circRNAs, but not intron-derived ciRNAs. **c,** Size distribution of circular RNAs, color-coded by ciRNAs (orange) and circRNAs (red), before and after filters. It showed that ciRNAs are relatively shorter than circRNAs and after above enrichment filtering, ciRNAs are mostly filtered out. **d**, circRNAs expressed in the *PICALM* locus in different cell types. **e**, Genes with more exons are generally more likely to produce more circRNAs. Exons being circularized are significantly longer than average. **f**, Scatterplot of circular reads vs. linear reads per junction sites (left) and per host gene (right). **g**, DisGeNet disease class enriched in the host genes of all detected circRNAs in this study. **h**, Percentage of synaptic genes among all circRNA host genes are significantly decreased in PD (t-test, $p = 0.036$) and in AD (t-test, $p = 0.012$) comparing their corresponding controls.

#### Supplementary Fig. 3

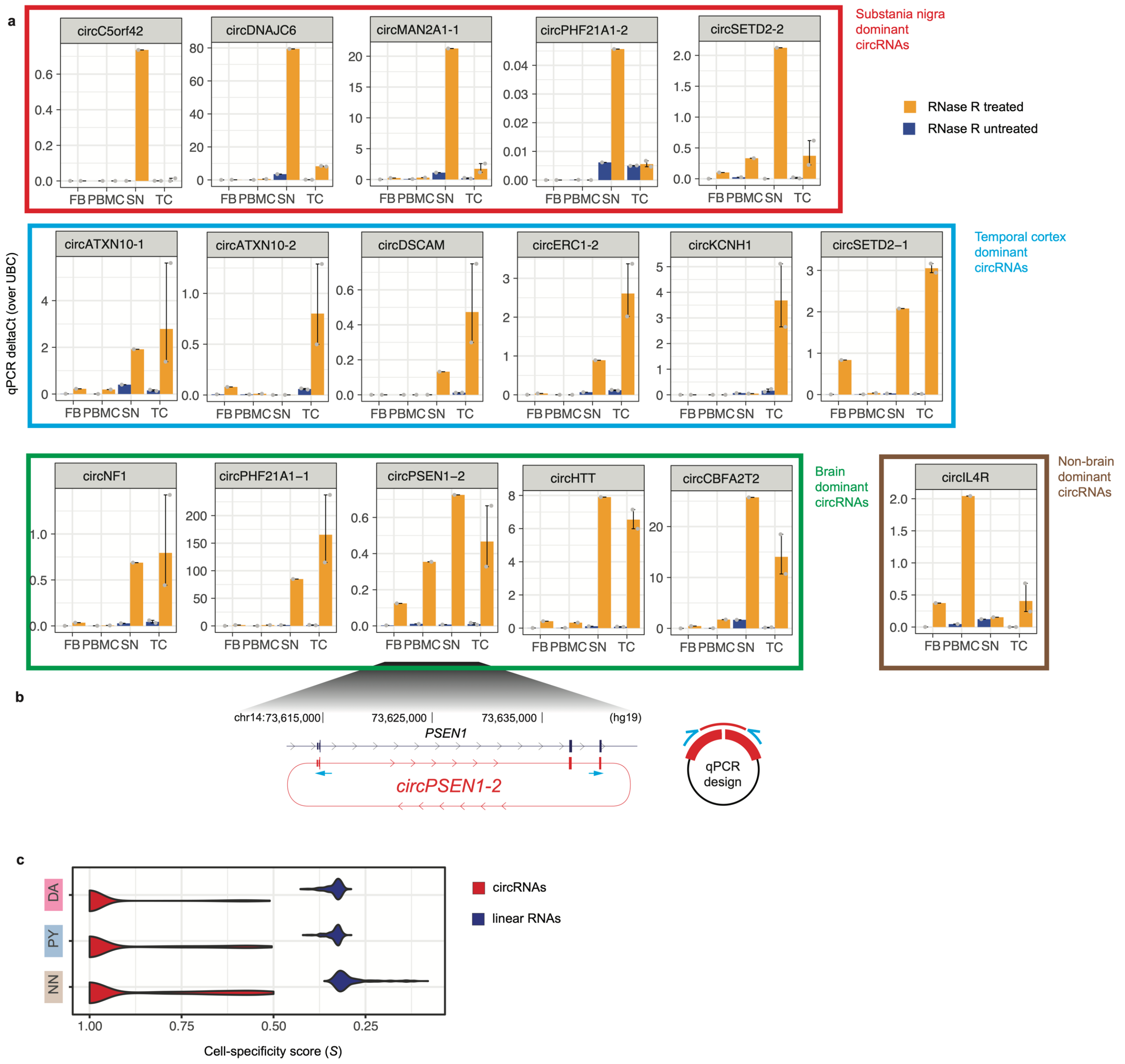

**Supplementary Figure 3. qPCR validation of circRNAs**. **a**, Barplot of qPCR delta Ct values (comparing to UBC control) for 17 select circRNAs, grouped into substantia nigro dominant circRNAs (in red frame), temporal cortex dominant circRNAs (blue frame), brain dominant circRNAs (green), and non-brain dominant circRNA (brown). The tissue dominance is based on visual inspection. Orange bars are for RNase R treated samples while darkblue bars are for untreated samples. Note that the qPCR validation were performed in bulk tissue (e.g. SN: susbtantia nigra and TC: temporal cortex), instead of laser-captured neurons, due to the requirement of input RNAs. **b,** Schema of qPCR design for circRNA validation using circPSEN1-2 as an example. Paired primers targeted the back-splicing junction sites were listed in Supplementary Table 4. **c**, Cell-specificity score distribution of circRNAs and linear mRNAs from the same genetic loci in three major cell types. This is limited to those host linear mRNAs which only produce one circular RNA.

#### Supplementary Fig. 4

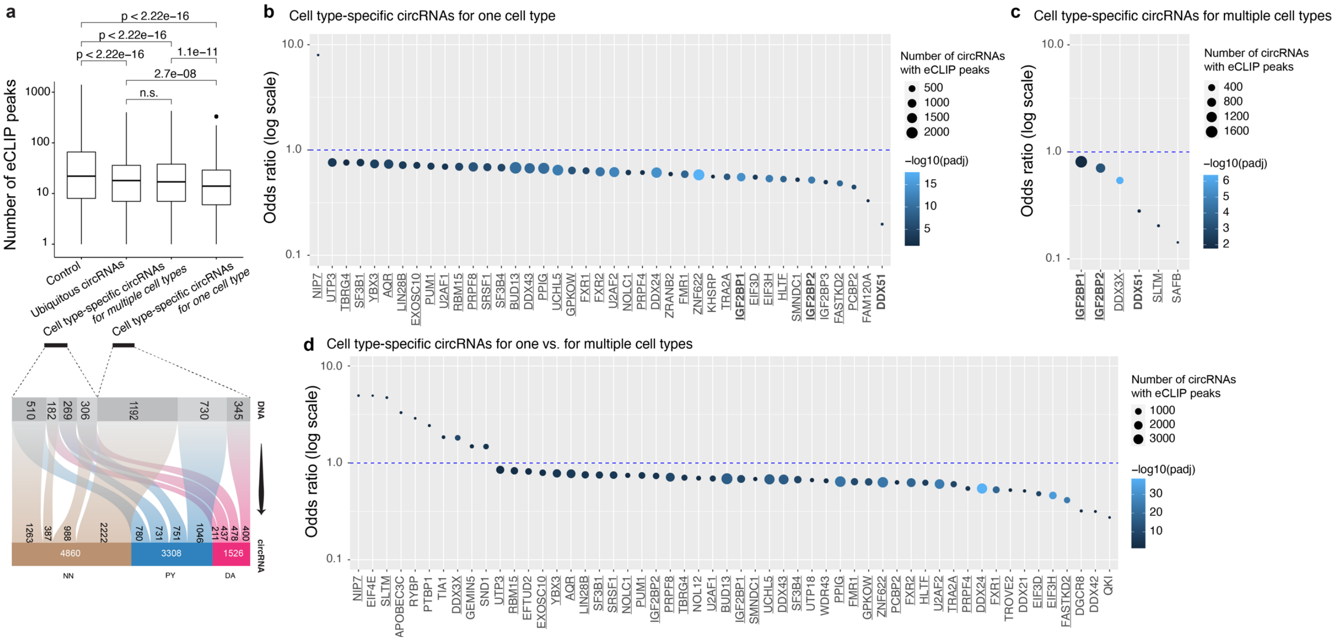


**Supplementary Fig. 4**. **RNA Binding Protein (RBP) activity and cell type-specific circRNAs. a**, Number of eCLIP peaks overlapping the two putative types of cell type specific circRNAs. **b**, Overlapped eCLIP binding peaks for 41 RBP were significantly positively (*NIP7*) or negatively enriched (all other RBPs) in circular exons of cell type-specific circRNAs for one cell type (e.g. circRNAs from locus producing *one or more cell type-specific* circRNAs specific to the *same one* cell type) compared to ubiquitous circRNAs with FDR < 0.05. Odds ratios from Fisher's exact test; $N = 5{,}608$ for circRNAs one cell type compared to $N = 1{,}942$ ubiquitous circRNAs. **c**. Overlapped eCLIP binding peaks for six RBPs were significantly less frequent at circular exons of cell type-specific circRNAs for multiple cell types (e.g., circRNAs from locus producing multiple cell type-specific circRNA for *multiple* cell types) compared to ubiquitous circRNAs. $N = 6{,}028$ circRNAs for cell type-specific circRNAs for multiple cell types compared to $N = 1{,}942$ ubiquitous circRNAs. **d**. Overlapped eCLIP binding peaks for 54 RBP were significantly positively ($N = 10$) or negatively enriched ($N = 44$) in circular exons of cell type-specific circRNAs *for one cell type* compared to cell type-specific circRNAs *for multiple cell types* with FDR < 0.05. Odds ratios from Fisher's exact test; $N = 5{,}608$ for circRNAs *for one cell type* compared to $N = 6{,}028$ circRNAs *for multiple cell types*.

#### Supplementary Fig. 5

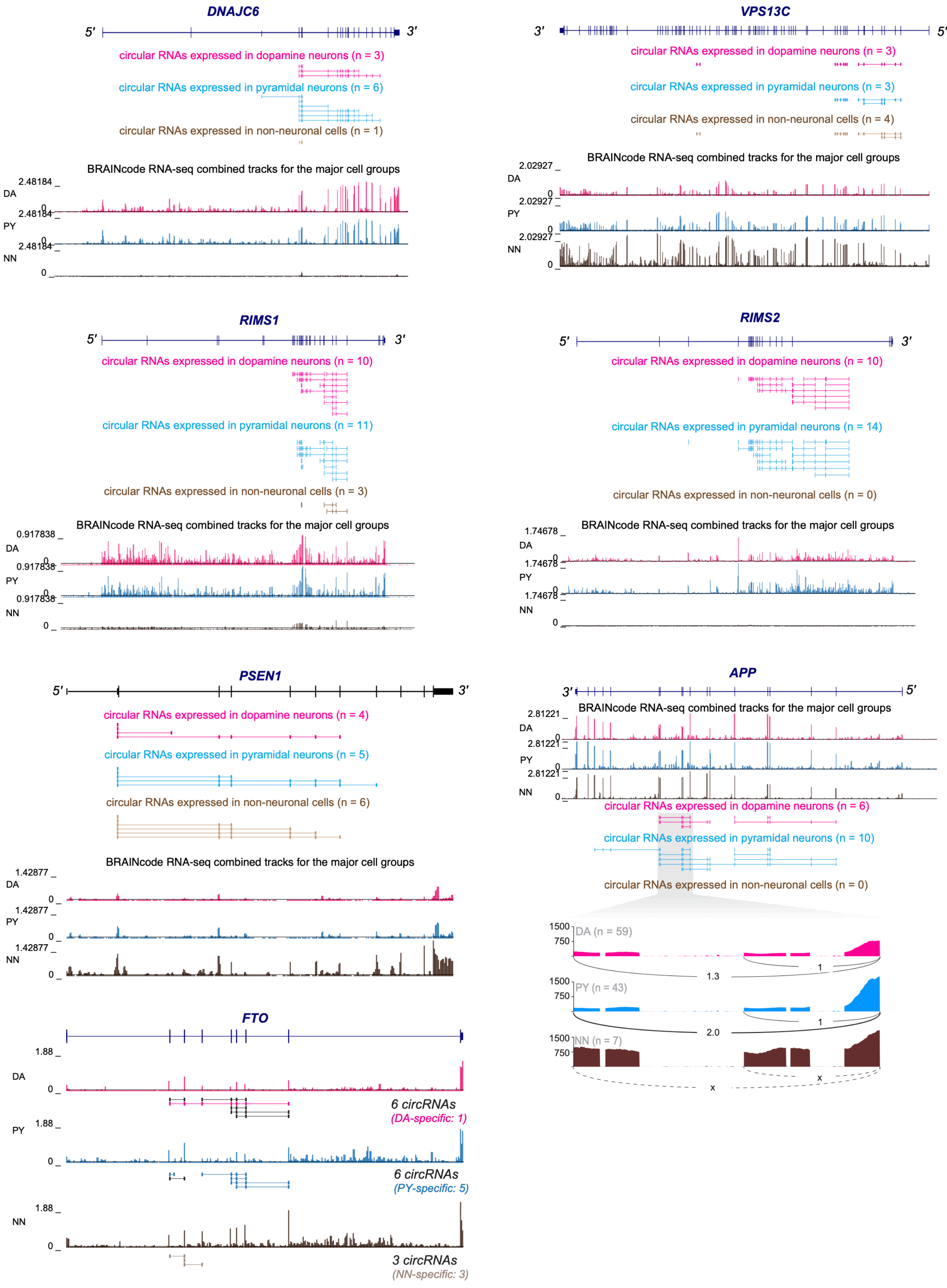
DNAJC6
5′
3′
circular RNAs expressed in dopamine neurons (n = 3)
circular RNAs expressed in pyramidal neurons (n = 6)
circular RNAs expressed in non-neuronal cells (n = 1)
BRAINcode RNA-seq combined tracks for the major cell groups
2.48184
DA
PY
NN
0
VPS13C
3′
5′
circular RNAs expressed in dopamine neurons (n = 3)
circular RNAs expressed in pyramidal neurons (n = 3)
circular RNAs expressed in non-neuronal cells (n = 4)
BRAINcode RNA-seq combined tracks for the major cell groups
2.02927
DA
PY
NN
0
RIMS1
5′
3′
circular RNAs expressed in dopamine neurons (n = 10)
circular RNAs expressed in pyramidal neurons (n = 11)
circular RNAs expressed in non-neuronal cells (n = 3)
BRAINcode RNA-seq combined tracks for the major cell groups
0.917838
DA
PY
NN
0
RIMS2
5′
3′
circular RNAs expressed in dopamine neurons (n = 10)
circular RNAs expressed in pyramidal neurons (n = 14)
circular RNAs expressed in non-neuronal cells (n = 0)
BRAINcode RNA-seq combined tracks for the major cell groups
1.74678
DA
PY
NN
0
PSEN1
5′
3′
circular RNAs expressed in dopamine neurons (n = 4)
circular RNAs expressed in pyramidal neurons (n = 5)
circular RNAs expressed in non-neuronal cells (n = 6)
BRAINcode RNA-seq combined tracks for the major cell groups
1.42877
DA
PY
NN
0
APP
3′
5′
BRAINcode RNA-seq combined tracks for the major cell groups
2.81221
DA
PY
NN
0
circular RNAs expressed in dopamine neurons (n = 6)
circular RNAs expressed in pyramidal neurons (n = 10)
circular RNAs expressed in non-neuronal cells (n = 0)
1500
750
DA (n = 59)
1.3
1
PY (n = 43)
2.0
1
NN (n = 7)
x
x
FTO
1.88
DA
0
6 circRNAs
(DA-specific: 1)
PY
6 circRNAs
(PY-specific: 5)
NN
3 circRNAs
(NN-specific: 3)

99

**Supplementary Figure 5. Locus plots for example circRNAs and their host loci,** including *DNAJC6, VPS13C, RIMS1, RIMS2, PSEN1,* and *APP*. Note that the cell-specific circRNAs are based on healthy samples only.

#### Supplementary Fig. 6

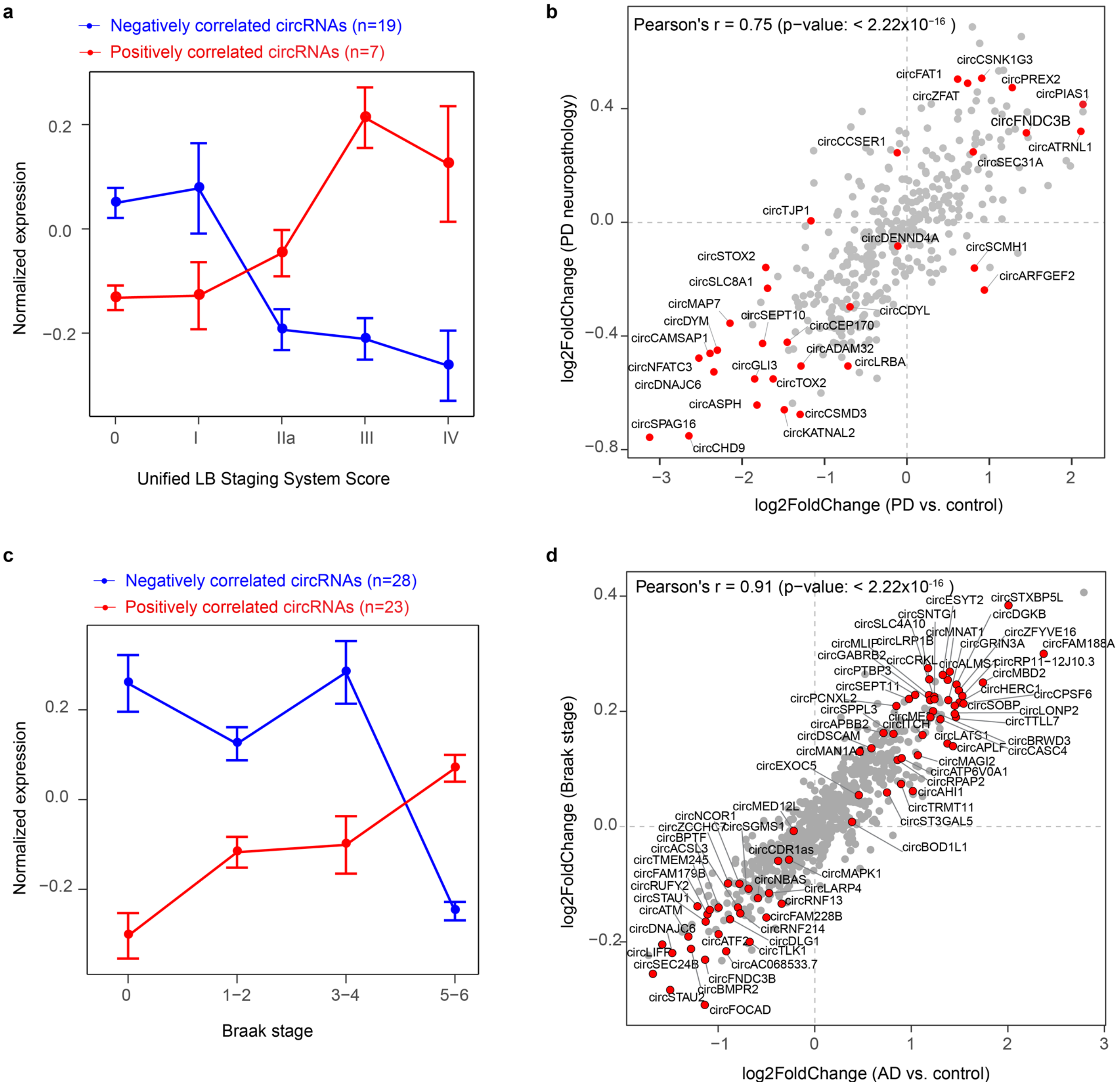


**Supplementary Figure 6. Exploring associations between circRNA expression and neuropathology.** **a**, Dopamine neuron circRNAs associated with Lewy body stages. The abundance of 26 circRNAs was associated with Lewy body stage with nominal $P$ values $\leq 0.05$ using linear regression analysis adjusted for covariates of sex, age, RIN, and PMI; 95 (out of 104) dopamine neuron transcriptomes with available neuropathology staging information (e.g., the Unified Lewy Body Staging System scores of 0, I, IIa, III, and IV, see Supplementary Table S1). None achieved the multiple-testing-corrected significance threshold of $FDR \leq 0.05$. The line graph shows mean and standard error of the normalized expression counts of positively and negatively associated circRNAs. **b**, Effect sizes of circRNAs associated with Lewy body neuropathology were highly correlated with the effect sizes for association with clinical diagnosis for the 18 samples with a clinical diagnosis of PD compared to 59 healthy controls without Lewy body neuropathology (Pearson's $r$ = 0.75, $P \leq 2.22\times10^{-16}$). Red dots, 32 suggestive circRNAs associated with Lewy body neuropathology or PD clinical diagnosis (e.g., $P < 0.05$ in either comparison); grey dots, circRNAs that were not associated with Lewy body neuropathology or PD clinical diagnosis or (e.g., $P \geq 0.05$ in both comparisons).

**c**, Pyramidal neuron circRNAs associated with AD neuropathology. The abundance of 51 circRNAs was associated with the neuropathological Braak stages of AD patients with nominal *P* values ≤ 0.05 using linear regression analysis adjusted for covariates of sex, age, RIN, and PMI ($N = 83$, including 9, 12, 11, 4, 4, 11, and 32 subjects with AD Braak stage of 0, 1, 2, 3, 4, 5, and 6, respectively; Fig. 1b). None achieved the multiple-testing-corrected significance threshold of $FDR < 0.05$. The line graph shows mean and standard error of the normalized expression counts of positively and negatively associated circRNAs.

**d**, Effect sizes of circRNAs associated with neuropathological AD Braak stages were highly correlated with the effect sizes for association with clinical diagnosis of AD compared to healthy controls (Pearson's $r = 0.91$; $P \leq 2.22 \times 10^{-16}$; $N = 43$ AD and 40 control samples; Fig. 1b). Red dots, 71 suggestive circRNAs associated with AD clinical diagnosis or AD neuropathology (e.g., $P < 0.05$ in either comparison); grey dots, circRNAs that are not associated with AD clinical diagnosis or AD neuropathology (e.g., $P \geq 0.05$ in both comparisons).
